# Inbreeding depression in an outbred nine-spined stickleback (*Pungitius pungitius*) population

**DOI:** 10.1101/2022.03.15.483967

**Authors:** Antoine Fraimout, Pasi Rastas, Lei Lv, Juha Merilä

## Abstract

Inbreeding depression refers to the reduced fitness of offspring produced by related individuals and is expected to be rare in large outbred populations. When it occurs, marked fitness loss is possible as large populations can carry large loads of recessive harmful mutations which are normally sheltered at the heterozygous state. Using experimental cross data and genome-wide identity-by-descent (IBD) relationships from an outbred marine nine-spined stickleback (*Pungitius pungitius*) population, we documented a significant decrease in offspring survival probability with increasing parental IBD sharing associated with an average inbreeding load (*B*) of 15.896. Interestingly, we found that this relationship was also underlined by a positive effect of paternal inbreeding coefficient on offspring survival, suggesting that certain combinations of parental inbreeding and genetic relatedness among mates may promote offspring survival. Apart from demonstrating substantial inbreeding load in an outbred population, the results also highlight the potential caveat associated with artificial establishment of families in experimental studies: wild founder individuals are often - and perhaps mistakenly - assumed to be unrelated.

## INTRODUCTION

Mating between close relatives often reduces the fitness of offspring through morphological, physiological or reproductive defects, a phenomenon known as inbreeding depression [1]. Inbreeding depression may lead in the most extreme case to offspring unviability and eventually to population extinction [2]. Quantitative and population genomic approaches to study inbreeding depression have become increasingly important as the environmental pressures imposed by global changes are predicted to increase habitat fragmentation, reduce population sizes and consequently increase the probability of inbreeding in the wild [3-5].

The magnitude of inbreeding depression depends on the interplay between the effective population size (*N*_*e*_), the rate at which new mutations arise in the population and the efficiency of selection at eliminating harmful mutations [6]. Populations with large *N*_*e*_ are large mutational targets but natural selection is more efficient against deleterious mutations in large than in small populations [7,8]. As a result, most mutations segregating in populations with high *N*_*e*_ are recessive, mildly deleterious, and can subsist in the population in the heterozygous state [9]. This in turn means that the cost of inbreeding in large populations is expected to be high, as it increases the chance of recessive mutations to be expressed as lethal homozygotes [10]. Conversely in small populations, and particularly those confined to population isolates with no or limited migration, the probability of mating with genetically related individuals is drastically increased. Nonetheless, the cost of inbreeding in such populations should be reduced, as deleterious mutations would be rapidly purged due to homozygotes lethality [6, 11, 12].

These theoretical expectations have been investigated in a variety of wild model species across different taxa [13, 1]. In a comprehensive empirical study of the link between *N*_*e*_ and genetic load in the water flea *Daphnia magna*, Lohr & Haag [14] reported a clear positive relationship between the load of segregating recessive mutations (expressed through inbreeding depression) and *N*_*e*_. Similarly, empirical work comparing large and small populations of *Arabidopsis thaliana* have shown that small-sized populations carry large genetic loads due to the fixation of deleterious mutations (*i*.*e*., drift load; [10]. To date, theoretical expectations regarding the relationship between *Ne*, genetic load and inbreeding depression have been verified in a number of terrestrial systems (*e*.*g*., [15-19])

In marine organisms, studies of genetic load are still rare [20] and have been often focused on specific groups (*e*.*g*., Oysters; [21-23]). The magnitude of inbreeding depression in fish species in particular seem to have been under-studied with the exception of classical models (*e*.*g*., guppies, [24, 25]; three-spined sticklebacks, [26, 27]; zebrafish, [28]) or species of economic interest (*e*.*g*., salmonids, [29, 30]). Yet, demographic and genetic parameters specific to marine organisms make them an opportune model to test theoretical predictions of inbreeding depression in the wild. Indeed, while several marine species are characterised by large *N*_*e*_, high dispersal and low genetic differentiation among populations [31-34], it has been long recognized that population structure in the sea may be more prevalent than previously thought [35-39] and that such structure can have major implications for evolutionary responses to selection [40, 41]. In this context, population genomics tools can shed light on the fine-scale relatedness patterns occurring in structured populations, and advance our understanding of inbreeding depression in wild outbred populations [42].

The main objective of this study was to investigate whether cryptic relatedness estimated from genome-wide identity-by-descent (IBD) relationships among wild-collected parents had an effect on offspring survival in a large crossing experiment of the nine-spined stickleback (*Pungitius pungitius*). Specifically, we tested if relatedness among a set of wild parental fish used to produce F_1_-generation sibs through artificial mating led to inbreeding depression in offspring viability. Our results show that genetic similarity among parents can yield substantial reduction in offspring survival even when the level of relatedness is low.

## MATERIAL AND METHODS

### Sampling and rearing

The fish used in this study came from a large crossing design used in previous studies [43]. Wild parental fish were caught from a marine population in the Baltic Sea in Finland (60°13’N, 25°11’E) and brought to the aquarium facility of the University of Helsinki. The crossing scheme consisted of a maternal half-sib design where 46 females were crossed to two different males resulting in a total of 86 half-sib/full-sib families (3 females were mated to a single male). Parental fish we crossed artificially following standard *in vitro* split-clutch approach for sticklebacks [44]. Eggs were stripped from gravid females by gently squeezing their abdomen and placed in two different petri dishes. Males were over-anesthetized using tricaine methanesulfonate (MS222) in order to dissect their testes, which were subsequently minced in the dish containing the eggs from their corresponding females. Fertilization was performed by mixing the eggs and sperm using a sterile plastic pipette and was ensured by looking for signs of post-fertilization membranes under a binocular microscope. Following egg hatching and after larvae started feeding, each half-sib family was thinned to approximately 25 F_1_ offspring per family and transferred to two large aquaria. Individuals were placed at random to ensure that half of each family was represented in each aquarium. Offspring were mass-reared and their parental identity were subsequently recovered from genotype data as explained in [43] (and see below). Offspring survival was scored as the number of F_1_ fish alive at the end of the experiment.

### Sequencing and genotyping

Parental fish were whole-genome sequenced (WGS) on an Illumina HiSeq platform (BGI, Hong-Kong) at 5-10X sequencing coverage and offspring were sequenced using the Diversity Arrays Technology (DarTseq technology, Pty Ltd, Australia). Fastq files were mapped to the contig assembly of the *P. pungitius* reference genome [45] using bwa-mem [46] and then sorted bams for each file were obtained using SAMtools [46]. We obtained genotype likelihoods using Lep-MAP3 [47]. The linkage mapping and the pedigree construction were conducted using Lep-MAP3. Offspring were assigned to parents using the Lep-MAP3’s IBD module by calculating the Mendelian genotypic error rate for each combination of parents and offspring and by taking all combinations with error rate < 10% (average error rate was 5.5%). Finally, offspring sex was obtained by comparing the sequencing coverage on X and Y specific regions using SAMtools depth. The final genomic dataset consisted of 49,493 biallelic SNPs.

### Estimation of genomic inbreeding and relatedness

We estimated the genomic inbreeding coefficient (*F*_*G*_) of all dams and sires from SNP data using the *--ibc* option implemented in the GCTA software (v.1.93.2, [48]). This metric provides a measure of inbreeding that is unbiased, as homozygote genotypes are scaled by their allele frequencies [48]. Second, we estimated the pairwise relatedness between each pair of parents by computing the Genomic Relationship Matrix (GRM) in GCTA using the *--make-grm* function. GCTA uses the allelic correlation coefficient (see [48] p. 76 and [49]) to construct the pairwise matrix of IBD coefficients among all individuals.

To ensure that our results were not influenced by the specific approaches used to estimate *F*_*G*_ and IBD in GCTA, we replicated all analyses based on alternative estimates using the software PLINK (v1.90, [50]). We used the *--het* option in PLINK to obtain individual inbreeding coefficients and the *--genome* option to estimate the proportion of genome shared IBD between all pairs of parents. Inbreeding coefficients estimated with PLINK correspond to excess homozygosity and deviation from Hardy-Weinberg equilibrium. In PLINK, the GRM is constructed following the approach of VanRaden [51] and based on the probability of IBD computed from identity-by-state values. We chose these two software as they provide two alternative approaches to the estimation of each parameter of interest, and have previously been shown to improve detection of inbreeding depression using genomic data [52] and particularly in population with large *Ne* [53]. For the sake of comparison, we follow the nomenclature of other studies (*e*.*g*., [42, 52, 54, 55]) and hereafter refer to GCTA estimates of inbreeding coefficient as *F*_*UNI*_ and PLINK estimates as *F*_*HOM*_. Similarly, estimates of relatedness using GCTA and PLINK are referred to IBD_Yang_ and IBD_VR_, respectively. Finally, to account for the possible effects of SNP filtering on the estimation of inbreeding and relatedness coefficients, all analyses were ran using estimates of *F*_*G*_ and IBD from data pruned for minimum allele frequencies (MAF < 0.05) and linkage disequilibrium (LD > 0.8) using the R package SNPrelate [56] or using unpruned data. Prior to all analyses, we removed all markers present on *P. pungitius* sex chromosome (Chromosome 12, [57]) to retain only autosomal SNPs.

### Statistical analyses

We ran generalized linear mixed models (GLMMs) to investigate the effect of parental relatedness and parental inbreeding on offspring survival using the *lme4* package (v.1.1-27.1; [58]) in R (v.4.1.1; [59]). Offspring survival was modelled as a binomial response variable using the *glmer* function in *lme4* using a logit link function. Fixed effect terms included parental relatedness (IBD; the proportion of genome shared IBD between each pair of parents), paternal inbreeding coefficient (*F*_SIRE_), maternal inbreeding coefficient (*F*_DAM_) and their two-way interactions. To account for the used maternal half-sib design, dam identity was used as a random effect in all models. Preliminary investigations of the data indicated that the inclusion of the dam random effect led to a change in the sign of the effect of parental relatedness on offspring survival, indicating distinct effects of within-family relatedness (relatedness among the triad of parents of each half-sib family) and between-family relatedness (relatedness across dams) on offspring survival. To account for these distinct effects, we used a technique of within-subject centering [60] by calculating the mean relatedness (*mR*) value of each half-sib family and added it as a fixed effect in our GLMMs. Similarly, we estimated the component of relatedness within half-sib families by normalizing individual values as *wR*=*IBD*-*mR*, for each pair of parents. Model selection was performed based on the Akaike Information Criterion corrected for small sample size (AICc; which balances model fit and complexity, [61]), by using the *dredge* functions in the R package *MuMin* (v.1.43.17, [62]). Collinearity among the predictor variables was assessed using the variance inflation factor (VIF) analysis by using the *vif* function in the R package *car* (v.3.0-11, [63]). Our final model had the following syntax:

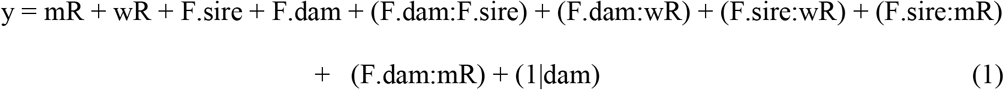

where *y* is the offspring survival coded as a two-column vector; colons indicate fixed terms interaction and (1|dam) corresponds to the notation of random effect of dam identity in the *lme4* syntax. We ran the model in equation (1) for all four combinations of SNP data filtering (pruned/unpruned) and genomic estimator (*F*_*UNI*_/*F*_*HOM*_) while keeping consistency between software used to estimate *F*_*G*_ and IBD (*i*.*e*., using *F*_*UNI*_ / IBD_Yang_ and *F*_*HOM*_ / IBD_VR_ together in the same models). Finally, as the use of a logarithmic link function to GLMMs has been advocated to ensure unbiased estimates of inbreeding load [55], we also ran all models using the “log” link option in *lme4* (see *Supplementary material*).

### Estimation of inbreeding load

Inbreeding depression can be quantified using the inbreeding load (*B*) expressing the number of lethal equivalents per gamete [64] and estimated as the negative slope of the regression of offspring inbreeding coefficients on the trait values on the logarithmic scale [55, 65]. Inbreeding coefficients of offspring are usually obtained from pedigree relationships which allow estimating the expected IBD relationships within individuals [66]. Here, as pedigree relationships would not capture any fine-scale variation of realized relatedness among parents (all parents are assumed to be unrelated), we estimated the magnitude of inbreeding load as the slope of the regression of parental relatedness (*mR*) on offspring survival from model (1) above. Finally, we calculated the mean and 95% confidence interval (CI) of *B* from 1000 bootstrap iterations for each combination of SNP data and genomic inbreeding estimator.

To ensure consistency of our results across statistical approaches and to make our results comparable to other studies, we also estimated *B* following the traditional approach using offspring inbreeding coefficients instead of parental relatedness as the main model’s fixed effect (see *Supplementary material*).

### Heritability estimates

We estimated the proportion of genetic variation explaining the variance if offspring survival by computing the heritability (*h*^*2*^) of survival rate using an animal model [13, 67] of the form:

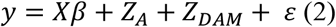

where *y* is the vector of offspring survival coded as a binary response variable (0 and 1 for dead and surviving individuals, respectively); *X* is the incidence matrix associated to the vector of fixed effect *β, Z*_*A*_ is the matrix of relatedness between individuals obtained from pedigree relationships; *Z*_*DAM*_ is the random effect of dam identity and *ε* is the error term.

We further tested if offspring survival variation resulted from inbreeding depression by adding individual *F*_*G*_ as a covariate in model (2). Indeed, inbreeding depression is expected to bias the estimation of the genetic variance components from animal models [68] and we hypothesized that a downward bias in *h*^*2*^ resulting from the added fixed effect of *F* would give further support to the role of inbreeding in survival rate variation in our data. Because genomic data to estimate offspring *F*_*G*_ was only available for surviving individuals, we assigned to each offspring the mean *F*_*G*_ value calculated from the survivors of each family (see also *Supplementary methods*). Animal models were fitted with the *MCMCglmm* R package [69] using the “threshold” family option appropriate for binomial regressions [70] and ran for 10,400,000 iterations with a burn-in period of 400,000 and sampling every 1000^th^ iteration. We ran standard model diagnostics for *MCMCglmm* models [69, 70] by investigating the trace plots of all variance components and computing their effective sample size using the *effectiveSize* function implemented in the *coda* R package [71] and checking for autocorrelations between iterations using the *autocorr* function from *coda*. We computed *h*^*2*^ both on the latent and observed scales using the *QGparams* function of the *QGglmm* R package [72].

## Results

### Inbreeding coefficients and relatedness among the parents

We found differences in inbreeding coefficients between sexes using both metrics, with males having lower inbreeding coefficients than females (*F*_*UNI*_: *t* = −5.497, *p* < 0.001; *F*_*HOM*_: *t* = −5.541, *p* < 0.001). Relatedness coefficients estimated with PLINK and GCTA were moderately positively correlated (*r* = 0.573; *p* < 0.001). Overall, relatedness between pairs of mates was low (Table S1), as expected from a random sample of individuals mated at random. The pairwise IBD estimated with PLINK ranged from 0 to 0.237 and from −0.029 to 0.090 with GCTA. Negative values estimated with GCTA indicate that these pairs of individuals were less related than the pairs in the sample on average [48].

### Inbreeding depression

There was a negative effect of parental relatedness on offspring survival across all models (Fig. 1, Table 1), but model complexity (*i*.*e*., inclusion of fixed terms and their interactions) varied depending on SNP data filtering and the estimator (*F*_*G*_ and IBD) used (Table 1). Using pruned SNP data, the full model using PLINK estimators (*F*_*HOM &*_ IBD_VR_) indicated a negative effect of parental relatedness on offspring survival (Table 1; *p* = 0.022) and that relatedness within half-sib families was positively associated with offspring survival (Table 1; *p* = 0.048). In other words, triads of parents that were on average more genetically related had fewer survivors among their progeny while females tended to have fitter offspring when mated with more related males. We also found contrasting effects of parental *F*_*HOM*_ on survival probability with more inbred females showing lower offspring survival (Table 1 *F*_*DAM*_; *p* = 0.022) but more inbred males showing higher offspring survival (Table 1, *F*_*SIRE*_; p < 0.001). We could not test for any further interaction terms due to a high degree of multicollinearity among fixed terms resulting in high VIF. Using unpruned SNP data, the model based on PLINK estimators yielded similar results with parental relatedness having a strong negative effect on offspring survival (Table 1; *p* < 0.001). Paternal inbreeding coefficient in this model was also found to be positively related to offspring survival (Table 1, *F*_*SIRE*_; *p* < 0.001). The relatedness within half-sib families had a negative effect on offspring survival in this model (Table 1, *wR*; *p* < 0.001).

**Table 1.** Results from the GLMMs on the effect of parental relatedness on offspring survival. For each model, the slope of regression for each coefficient of the fixed terms and interactions is given along with the standard error (SE) and the *p*-value. *mR:* mean relatedness between pairs of parents; *wR* normalized within-half-sib relatedness; *F*_*DAM*_: inbreeding coefficient of the dam; *F*_*SIRE*_: inbreeding coefficient of the sire. Colons indicate interactions between terms. Statistically significant p-values (*p* < 0.05) in bold.

| SNP data | IBD Estimator | $F_G$ Estimator | Fixed effects and interactions | Slope | SE | $p$-value |
| --- | --- | --- | --- | --- | --- | --- |
| Pruned | IBD<sub>VR</sub> | $F_{HOM}$ | mR | -6.797 | 2.979 | **0.022** |
|  |  |  | wR | 3.465 | 1.759 | **0.048** |
| | | | $F_{DAM}$ | -7.180 | 3.147 | **0.022** |
| | | | $F_{SIRE}$ | 9.828 | 1.479 | **< 0.001** |
| | IBD<sub>Yang</sub> | $F_{UNI}$ | mR | -19.380 | 6.5040 | **0.003** |
|  |  |  | wR | -7.940 | 5.752 | *0.167* |
| | | | $F_{DAM}$ | -15.446 | 4.961 | **0.002** |
| | | | $F_{SIRE}$ | 14.955 | 2.177 | **< 0.001** |
| | | | $F_{DAM}:F_{SIRE}$ | -195.637 | 64.489 | **0.002** |
| | | | wR:$F_{DAM}$ | 475.043 | 201.783 | **0.018** |
| Unpruned | IBD<sub>VR</sub> | $F_{HOM}$ | mR | -12.563 | 1.640 | **< 0.001** |
|  |  |  | wR | -9.978 | 2.571 | **< 0.001** |
| | | | $F_{SIRE}$ | 20.966 | 2.761 | **< 0.001** |
| | IBD<sub>Yang</sub> | $F_{UNI}$ | mR | -25.334 | 9.494 | **0.008** |
|  |  |  | wR | 1.8537 | 5.909 | *0.754* |
| | | | $F_{SIRE}$ | 18.095 | 3.584 | **< 0.001** |

**Figure 1.**
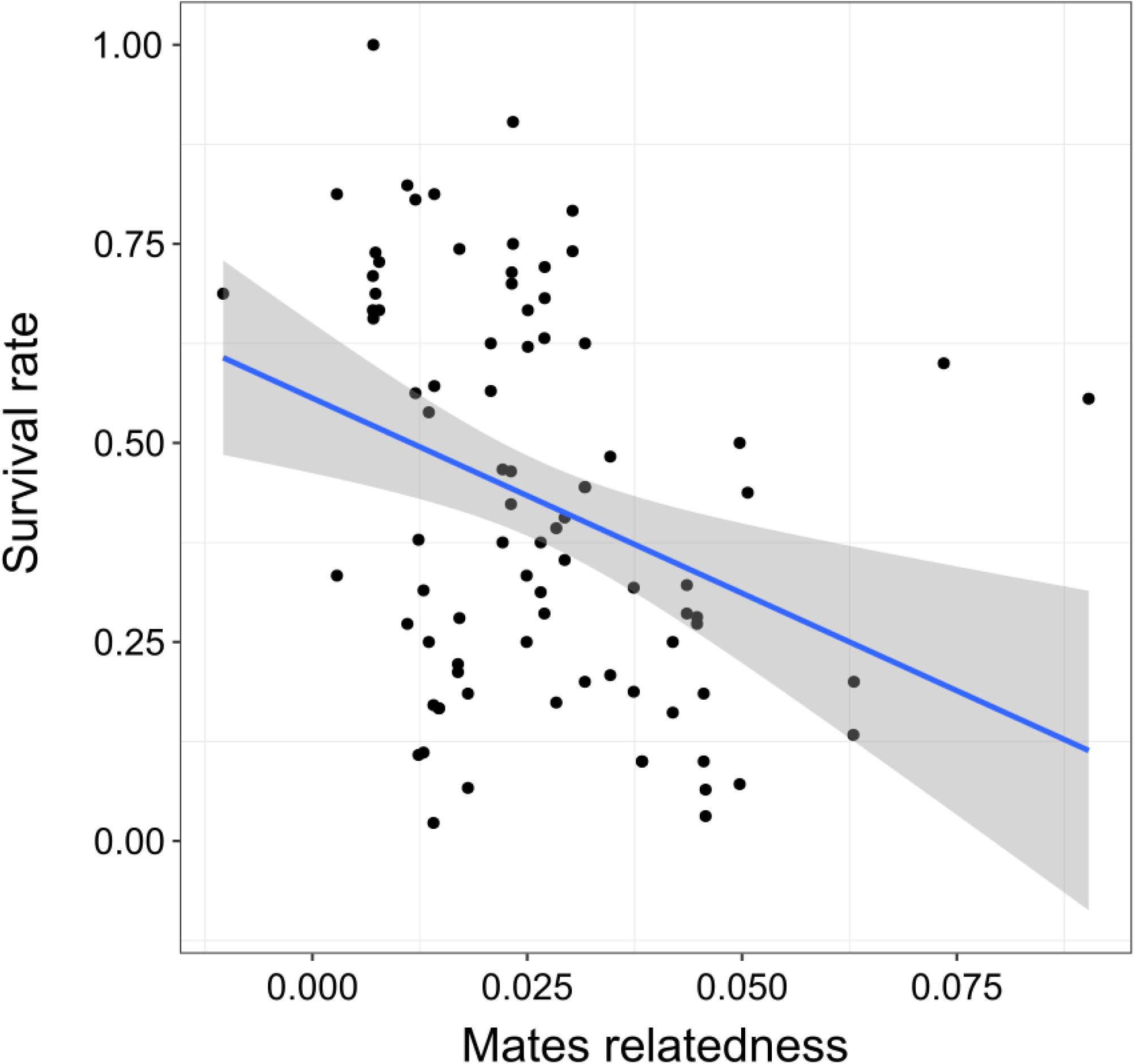
Inbreeding depression in juvenile survival rate of nine-spined sticklebacks. The survival rate for each family (black filled circles) is shown as a function of the relatedness coefficients between pairs of mates estimated with GCTA (see *Methods*). Blue line represents the fitted regression line from the ggplot *smooth* function and grey shadings are the 95% confidence intervals around the predicted values.

Models based on the GCTA estimators (*F*_*UNI &*_ IBD_Yang_) also indicated a negative effect of parental relatedness on offspring survival (Fig. 1, Table 1). When using pruned SNP data, we found again opposing effects of parental inbreeding coefficients on offspring survival (Table 1). This was also reflected by the significant negative interaction between the *F*_*DAM*_ and *F*_*SIRE*_ terms in the model (Table 1, *p* = 0.002). Finally, the GLMM based on unpruned SNP data showed a negative effect of parental relatedness on offspring survival (Table 1, *p* = 0.008) and a positive relationship between paternal inbreeding coefficient and survival (Table 1, *p* < 0.001).

Overall, estimates of *B* varied depending on the type of filtering and genomic predictor used, and showed relatively large confidence intervals around the mean (Table 2). The mean *B* across all models was 15.896. Results based on the clutch-averaged inbreeding coefficients (see *Supplementary methods*) were concordant with the above estimates with a mean *B* = 16.946 across models (Table S2). Finally, results obtained using the log-link function gave highly similar results for all GLMMs (Table S3).

**Table 2.** Bootstrap estimates of the inbreeding load. The mean values for the inbreeding load (*B*) are reported along with their 95% Confidence Intervals (CI). The type of SNP data, IBD and *F*_*G*_ estimators are as defined in the main text.

| SNP data | IBD Estimator | $F_G$ Estimator | B | 95% CI |
| --- | --- | --- | --- | --- |
| Pruned | IBD<sub>Yang</sub> | $F_{UNI}$ | 19.332 | [7.430 - 31.274] |
| Unpruned | IBD<sub>Yang</sub> | $F_{UNI}$ | 24.847 | [9.243 - 42.119] |
| Pruned | IBD<sub>VR</sub> | $F_{HOM}$ | 7.039 | [2.974 - 12.386] |
| Unpruned | IBD<sub>VR</sub> | $F_{HOM}$ | 12.367 | [9.529 - 15.658] |

### Heritability of offspring survival

We found moderate *h*^*2*^ in offspring survival with *h*^*2*^ = 0.17 [0.04;0.32] on the latent scale and *h*^*2*^ = 0.13 [0.04; 0.25] on the observed scale. Adding offspring inbreeding coefficients as covariate in the animal model reduced estimates of *h*^*2*^ on both scales with *h*^*2*^ = 0.05 [1.7e-09; 0.12] and *h*^*2*^ = 0.06 [2.12e-09; 0.14] on the latent and observed scales, respectively.

## Discussion

The most salient finding of this study was the fairly severe inbreeding load observed across all GLMMs, seemingly much more severe than in other taxa for juvenile survival. This result shows that large outbred populations can harbour substantial inbreeding load, and more importantly, that inbreeding depression can ensue even when levels of parental relatedness is low. In the following, we discuss the implications of these results for the study of inbreeding depression in the wild.

The magnitude of observed inbreeding depression exceeds that of a wide array of taxa previously reported in the literature. Early estimates of lethal equivalents in several mammals and bird species were on average one order of magnitude lower for survival traits ([13], see their Table 1 p. 279) compared to our estimates. Recent surveys of the literature [55] are in line with this observation and in the particular case of juvenile survival, inbreeding loads rarely exceed 1-3 homozygote lethals. Although our estimates present relatively wide confidence intervals and vary among the genomic estimators used, the order of magnitude and range of the inbreeding loads reported here is on par with severe loads previously reported from some natural populations [73-76]. This has important conservation implications [77] as the results suggest that large-sized populations may be at a high risk of extinctions if rapid decline in population size or habitat fragmentation would increase the probability of inbred mating.

The severity of inbreeding depression is known to depend on the environmental conditions under which it is expressed, with stressful conditions amplifying its severity [78, 79]. Since our study was carried out in a laboratory, we may have underestimated its real magnitude. Indeed, wild *P. pungitius* encounter parasites [80], predators and other severe abiotic challenges which could potentially amplify the negative consequences of inbreeding. Conversely, it is difficult to dismiss the possibility that the unnatural laboratory environment could also lead to overestimation of the magnitude of inbreeding depression, although this possibility seems unlikely. Regardless of the ‘true’ natural level of inbreeding depression, the results show potential for it to exist in this outbred marine population.

An unexpected and intriguing result was the positive effect of paternal inbreeding on offspring survival: offspring sired by more inbred males showed increased survival probability, regardless of the estimator of inbreeding coefficient used. Given the negative relationship between parental relatedness and offspring survival, it is possible that inbred males would benefit from mating with more distantly related females. Similar results have been reported in the gopher tortoise (*Gopherus polyphemus*) in which increased maternal inbreeding was associated with higher offspring fitness despite a strong negative effect of parental relatedness [82]. In their study, Yuan *et al*. [82] propose that such mating strategy (kin avoidance by inbred individuals) would alleviate the fitness cost of inbreeding, particularly in the sex investing more resources in reproduction. In *P. pungitius*, males provide parental care through nest-building, guarding and fanning of eggs and could thus benefit from this mating strategy. Unfortunately, due to multicollinearity among variables, our current data did not allow us to model the interaction between paternal inbreeding and parental relatedness and test this hypothesis further. However, we found that survival rate was heritable, indicating that genetic differences between families explain some amount of variation in offspring viability. Hence, certain combinations of parental inbreeding coefficients and genetic relatedness yielding higher survival could be favoured by selection and provide basis to the evolution of mate choice in this species (cf. [83, 84]). However, heritability of offspring viability was lowered substantially when accounting for individual inbreeding coefficients, suggesting that inbreeding affected survival to a larger extent than among-family genetic differences. Experimental studies of mate choice controlling for the effect of parental inbreeding coefficient and parental relatedness should be particularly interesting to address these questions.

While the results demonstrate that potential for inbreeding depression is present in the study population, the results should not be extrapolated to say that it would be realized in wild *P. pungitius* populations. This because all mating in this study were artificial, and therefore eluded all aspects of mate choice, including kin recognition and active inbreeding avoidance. Such behaviours have been studied in the related three-spined stickleback (*Gasterosteus aculeatus*) in which gravid females prefer to lay eggs in nests built by unrelated males [85]. However, such studies of mate choice and inbreeding avoidance often use very different levels of relatedness by contrasting completely unrelated to relatively closely related individuals (*e*.*g*., full-sib). In the present study, the range of relatedness among parental fish was narrow, and well below full-sib relationship (the maximum value of IBD_VR_ = 0.237 corresponding to mating with aunt/uncle or half-sib). Hence, it is unclear whether kin recognition in sticklebacks would be fine-tuned to the levels of relatedness measured in our study and whether wild females could discern between males with such low levels of relatedness. Furthermore, high levels of local kinship have been reported in other marine fish [39, 86] and the relatively strong population structure found in marine populations of *P. pungitius* [41, 87] could promote similar local kinship patterns and increase the probability of inbred mating despite large population sizes.

In conclusion, as predicted by theory, the results show that there is an opportunity for severe inbreeding depression to occur in large outbreed marine populations. Future studies of inbreeding depression in the wild should benefit from the inclusion of fine-scale measurement of parental relatedness in models of inbreeding depression.

## Ethical statement

This experiment was approved by the National Animal Experiment Board, Finland (permission numbers: ESLH-STSTH223A and STH037A)

## Data accessibility statement

Sequence data that support the findings of this study have been deposited in European Nucleotide Archive (ENA) under the Accession nos PRJEB39736 (linkage map parents), PRJEB39760 (linkage map offspring).

## Acknowledgments

We thank everyone who helped with catching and rearing the fish, and Takahito Shikano, Chris Eberlein, and Sami Karja in particular. Thanks are also due to Laura Hänninen, Miinastiina Isssakainen, Sami Karja and Kirsi Kähkönen for sample preparations and DNA extractions. Ari Löytynoja and Mikko Kivikoski are thanked for access to genotype data. Our research was supported by the Academy of Finland grants (# 129662, 134728 and 218343) to JM. The computing resource support from CSC - the Finnish IT Center for Science Ltd administered by the Ministry of Education and Culture, Finland is gratefully acknowledged.

