## supplementary material for "Inbreeding depression in an outbred nine-spined stickleback (*Pungitius pungitius*) population"

Antoine Fraimout*[1,2] Pasi Rastas[3], Lei Lv[2,4] and Juha Merilä[1,2]

[1]Ecological Genetics Research Unit, Organismal and Evolutionary Biology Research Programme, Faculty of Biological and Environmental Sciences, FI-00014 University of Helsinki, Finland
[2]Area of Ecology and Biodiversity, School of Biological Sciences, The University of Hong Kong, Hong Kong SAR
[3]Institute of Biotechnology, University of Helsinki, Finland
[4] School of Ecology, Sun Yat-Sen University, Guangzhou, China

### Table of contents


| | |
|---|---|
| **Supplementary methods** | Page 2 |
| **Figure S1. Approaches to estimate inbreeding depression**. | Page 3 |
| **Table S1. Estimates of genomic inbreeding coefficient and relatedness among parental fish.** | Page 4 |
| **Table S2. Bootstrap estimates of the inbreeding load using clutch-averaged $F_G$.** | Page 5 |
| **Table S3. Results from the GLMMs on the effect of parental relatedness on offspring survival (log-link).** | Page 6 |

***Supplementary methods***

*Estimation of inbreeding load from offspring inbreeding coefficients*

Inbreeding depression is traditionally estimated from the regression of inbred offspring's fitness (the fitness measure of offspring whose parents are related) on their inbreeding coefficient (*F*) [1]. Here we used the genomic inbreeding coefficient ($F_G$) calculated from sequence data to estimate each individual's inbreeding in our data. Because only surviving individuals were genotyped at the end of the experiment, we estimated inbreeding depression as the inbreeding load (*B*) corresponding to the negative slope of the regression of parental relatedness on the survival rate of their offspring (Fig. S1A; and see *Main text*).

We also used an alternative approach of estimation where each offspring within a family was assigned with the within-clutch average value of $F_G$ estimated from its surviving full-siblings (*Clutch average* $F_G$ in Fig. S1B). To this end, we ran generalized linear mixed models (GLMMs) to investigate the effect of offspring inbreeding coefficients on survival using the *lme4* package (v.1.1-27.1; [2]) in R (v.4.1.1; [3]). Offspring survival was modelled as a binary response variable using the *glmer* function in *lme4*. Contrary to the analyses described in the main text, we modelled offspring survival using a poisson distribution by setting the *family* option of the *glmer* function to *poisson.* We ran the following GLMM:

$$y = \text{F.offspring} + \text{F.sire} + \text{F.dam} + (1|\text{dam}) \qquad (1)$$

where *y* is the offspring survival coded as a binary (0: dead, 1: alive) vector; *F.offspring* is the average clutch $F_G$ and (1|dam) corresponds to the notation of random effect of dam identity in the *lme4* syntax. We ran the model in equation (1) for all four combinations of SNP data filtering (pruned/unpruned) and genomic estimator ($F_{UNI}$/$F_{HOM}$) while keeping consistency between software used to estimate $F_G$ and IBD (*i.e.*, using $F_{UNI}$ / $\text{IBD}_{\text{Yang}}$ and $F_{HOM}$ / $\text{IBD}_{\text{VR}}$ together in the same models).

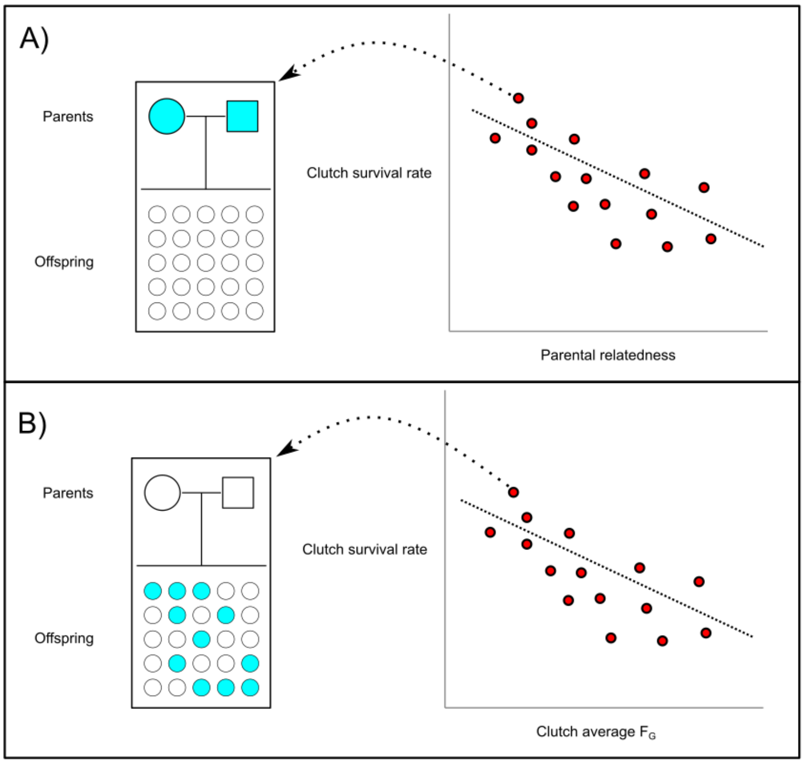


**Figure S1. Approaches to estimate inbreeding depression**. A) Inbreeding depression is estimated as the slope of the regression of offspring survival (y axis) on parental relatedness (x axis). B) Inbreeding depression is estimated as the slope of the regression of offspring survival (y axis) on the average inbreeding coefficient of the clutch. Blue dots represent the individuals used to estimate the parameter of interest (IBD or $F_G$), either the parents (A) or the survivors from each clutch (B); red dots represent each family.

**Table S1. Estimates of genomic inbreeding coefficient and relatedness among parental fish.** Individual inbreeding coefficients ($F_G$) were estimated using the $F_{UNI}$ metric implemented in GCTA or $F_{HOM}$ using PLINK. Relatedness between parents ($IBD_{pair}$) was estimated from the GRM constructed with GCTA or PLINK. For each metric, mean and standard deviation (SD) are reported.

| **Parameter** | **Metric** | **Sex** | **Mean (SD)** |
|---|---|---|---|
| $F_G$ | $F_{UNI}$ (GCTA) | Males | -0.042 (0.043) |
| | | Females | -0.001 (0.034) |
| | $F_{HOM}$ (PLINK) | Males | 0.335 (0.057) |
| | | Females | 0.389 (0.044) |
| $IBD_{pair}$ | GRM (GCTA) | Both | 0.026 (0.021) |
| | GRM (PLINK) | Both | 0.066 (0.054) |

**Table S2. Bootstrap estimates of the inbreeding load using clutch-averaged $F_G$.** The mean values for the inbreeding load (B) are reported along with their 95% Confidence Intervals (CI). The type of SNP data, IBD and $F_G$ estimators are as defined in the main text.

| SNP data | IBD Estimator | $F_G$ Estimator | B | 95% CI |
|---|---|---|---|---|
| Pruned | $IBD_{Yang}$ | $F_{UNI}$ | 14.310 | [12.141 - 16.565] |
| Unpruned | $IBD_{Yang}$ | $F_{UNI}$ | 21.706 | [17.956 - 25.627] |
| Pruned | $IBD_{VR}$ | $F_{HOM}$ | 12.410 | [10.122 - 14.676] |
| Unpruned | $IBD_{VR}$ | $F_{HOM}$ | 19.360 | [16.132 - 22.614] |

**Table S3. Results from the GLMMs on the effect of parental relatedness on offspring survival (log-link).** For each model, the slope of regression for each coefficient of the fixed terms and interactions is given along with the standard error (SE) and the *p*-value. *mR:* mean relatedness between pairs of parents; *wR* normalized within-half-sib relatedness; $F_{DAM}$: inbreeding coefficient of the dam; $F_{SIRE}$: inbreeding coefficient of the sire. Colons indicate interactions between terms. Statistically significant p-values ($p < 0.05$) in bold.

| SNP data | IBD Estimator | $F_G$ Estimator | Fixed effects and interactions | Slope | SE | *p*-value |
|---|---|---|---|---|---|---|
| Pruned | $IBD_{VR}$ | $F_{HOM}$ | mR | -3.900 | 1.662 | **0.019** |
| | | | wR | 1.654 | 1.001 | *0.098* |
| | | | $F_{DAM}$ | -3.878 | 1.769 | **0.028** |
| | | | $F_{SIRE}$ | 6.696 | 0.982 | < **0.001** |
| | $IBD_{Yang}$ | $F_{UNI}$ | mR | -11.0115 | 3.688 | **0.003** |
| | | | wR | -2.123 | 2.795 | *0.447* |
| | | | $F_{DAM}$ | -7.933 | 2.758 | **0.004** |
| | | | $F_{SIRE}$ | 9.288 | 1.303 | < **0.001** |
| | | | $F_{DAM}$:$F_{SIRE}$ | -121.525 | 39.831 | **0.002** |
| | | | wR:$F_{DAM}$ | 290.543 | 112.701 | **0.009** |
| Unpruned | $IBD_{VR}$ | $F_{HOM}$ | mR | -7.303 | 0.856 | < **0.001** |
| | | | wR | -6.398 | 1.429 | < **0.001** |
| | | | $F_{SIRE}$ | 14.517 | 1.679 | < **0.001** |
| | $IBD_{Yang}$ | $F_{UNI}$ | mR | -14.673 | 5.526 | **0.008** |
| | | | wR | 3.457 | 2.964 | *0.244* |
| | | | $F_{SIRE}$ | 12.414 | 2.266 | < **0.001** |
